# When your border and my border differ: Spatial regionalisation and route choice depends on perceived landmark categorization

**DOI:** 10.1101/2022.11.10.515979

**Authors:** Lilian LeVinh, Hanspeter A. Mallot

## Abstract

Humans spontaneously structure spatial environments into a hierarchy of regions. Besides connectivity and boundaries, sensory or functional similarities of landmarks have been shown to influence construction of regions. In natural spaces, regions and places are usually named and these names provide additional cues for the cognitive construction of regions. In this study we investigate the role of semantic similarities of place names in the formation of regions in a virtual environment with homogeneous connectivity. Place names were selected and distributed in a way that two alternative semantic categorizations were possible which corresponded to two equally valid regionalisations of the same environment. Region perception was assessed by having subjects choose between two equidistant routes crossing different numbers of region boundaries (classification consistent route choice). In a priming phase, subjects were biased to sort the landmark names in one or the other categorization scheme. In the test phase, subjects preferably selected routes with a lower number of region crossings according to the categorization previously discovered. The results show that perceived semantic similarity of place names does affect route choice and the formation of spatial regions.

## 1 Introduction

### 1.1 Regionalisation and Hierarchy

Imagine going to the zoo, a child points at an antelope and asks ‘What is that?’. The answer could simply be ‘An antelope’. The answer could also be on a more superordinate level such as ‘mammal’ or even ‘animal’. An answer focusing on subordinate level information is also possible, such as ‘This is Gertrude, she is 5 years old and has a slight limp on her left rear foot’. However, it is quite likely, that only the first answer will be seen as helpful, as the super- and subordinate level answers may be too abstract or too specific respectively. As can be seen from the example, objects can be represented on several hierarchical levels, which carry different kinds of information and vary in accessibility (Rosch et al., 1976). Hierarchical structures in representations lead to higher efficiency in information storage and planning (Cohen, 2000; Norman, 1981; Tomov et al., 2020). By placing concepts in superordinate groups, humans can avoid explicitly storing facts and attributes separately for each member of the group. Thus, if an attribute is characteristic of a group, by simply inferring that a member has that characteristic one avoids having to explicitly link the characteristic with each member. However, it can also lead to lower precision (Norman, 1981), for instance inheritance can also lead to wrong assumptions if a member of the group does not share a trait typical to the superordinate group (Stevens & Coupe, 1978). In cognitive representations, hierarchical structure is ubiquitous with respect to not only object and social categorization but also in the domain of space and action control (Huys et al., 2015). Experimental results suggest that spatial information is stored hierarchically as well (Hirtle & Jonides, 1985; McNamara et al., 1989). Both studies used an ordered tree algorithm to identify clusters within recall data. This was supported by subjects typically overestimating distances between objects the algorithm deemed from different groups, compared to within distances.

Hierarchical control is the regulation of action by superordinate and subordinate goals or plans. Superordinate and subordinate goals create a chain of behaviours which ultimately leads to the completion of a defined task. A superordinate goal contains several subordinate goals and does not have to resemble the subordinate goals (Logan & Crump, 2011). The superordinate goals are broader and more abstract, and can stay fixed, even if the subordinate goals change. For example, if the superordinate goal is to eat out, it is still maintained regardless of whether one decides for Italian or Japanese food. In turn, the goal of reaching the designated restaurant is achieved regardless of whether one goes by bike or car. One example where representational and control hierarchies interact is route planning in complex environments, route choices depend on regional knowledge and generate control chains which may themselves be hierarchically structured. Route choices can therefore be used to infer the regionalisation of spatial memories. In Wiener & Mallot (2003), Balaguer et al. (2016), Tomov et al. (2020), as well as in our study there is an interplay of both control and representational hierarchy. While route biases in our experiment are based on the regions in the control hierarchy, it would be the representational hierarchy that shapes the regions. While the existence of different hierarchical levels of planning does not seem to be controversial, the question of which higher level of entities form the basis of these hierarchies is largely unsolved.

Grouping criteria are diverse and not deterministic and there are many factors that can elicit regionalisation. An important factor would be similarity of the subordinate entities such as places or smaller regions, that are coerced into a larger region (Balaguer et al., 2016; Wiener & Mallot, 2003). While visual similarity may be the most apparent, remembered attributes and properties of geographical places can also be used in formation of regions (Friedman & Brown, 2000), where subjects formed regions encompassing several areas even from different countries and may have followed climate in doing so. Another regionalisation factor would be the structure of the environment, and physical borders encompassing an area have been found to play a strong role in compartmentalizing the environment (He & Brown, 2019; Meilinger et al., 2016). Designated borders, such as political borders (Stevens & Coupe, 1978), do not have to be clearly visible or follow the structure of the environment and thus may have to be learned. However, administrative boundaries such as national borders have also been shown to lead to regionalisation (Carbon & Leder, 2005; Uttal et al., 2010). Schick et al. (2019) have also found functionality as a factor. Grouping based on functionality was defined as grouping based on the relevance to a behavioural task.

In natural environments, cues or landmark traits are rarely as distinct as they are in experiments. For instance, some location may not show traits typical of their region, a skyscraper among two story buildings for example, or enclosing boundaries may be interrupted at several places. In addition, natural environments usually are more cluttered than the controlled lab environments and different people may thus focus on different cues within the environment (Griesbauer et al., 2022). Another reason why a region may be hard to pinpoint is that the regionalisation or grouping of items in general does not have to be based on well-defined borders. Regionalisation also has been shown to occur even if no groups or explicit grouping criteria were presented. Thus, grouping can influence task performance even if the grouping is based on features unrelated to the actual task (Tomov et al., 2020; Vickery & Jiang, 2009; Wiener & Mallot, 2003). However, the criteria are not a guarantee for regionalisation and and cues might not even be cummulative.

Another factor which complicating predictability for regionalisation is, that both region perception and the resulting biases can be modulated by context. With information that a place may not follow the superordinate region’s trend people may re-assess their judgements for example. Thus people may separate the same environment into different regions depending on the context.

#### Regions can influence the perception of space

According to Couclelis’ anchor point hypothesis (Couclelis et al., 1987), the cognitive map is organized hierarchically, with hierarchies defined by anchor points. Anchor points are thus named as they anchor elements within a region to a local reference frame. Unlike landmarks which are perceptually defined, anchor points are more personal and are more specific to an individual’s cognitive map. For instance, someone’s home as an anchor point has special meaning only to them. Anchor points are one step up in the formation of hierarchical information and abstract spatial concepts. Therefore, anchor points can influence associated elements due to the inheritance of the anchor’s attributes to the associated elements. Thus, if for example new information reveals a location to be more north than expected, subjects would also shift places of the same region northward to preserve their relative distances (Friedman & Brown, 2000; Friedman & Montello, 2006). Couclelis et al. (1987) describes this occurence as the tectonic plate hypothesis which is based on objects within a region being more closely linked with their anchor point than they are with elements outside the region.

The misattribution of the superordinate group traits to the group’s members can affect spatial cognition outside of anchor points (Friedman & Brown, 2000; Friedman & Montello, 2006; Stevens & Coupe, 1978; Tversky, 1992) Friedman & Montello (2006) found for instance, that even relative experts make systematic errors in the judgement of city locations: when estimating the latitudes of the cities, the estimates followed the respective countries’ relative positions. For example, while the Canadian city of Toronto is south of Seattle (USA), subjects tended to place Toronto up north, and vice versa for the Mexican city Ciudad Juárez, which is north to some US cities such as Houston. One of the best known examples of categorical errors in spatial reasoning is the Miami-Lima illusion (Stevens & Coupe, 1978; Tversky, 1981) describing the tendency of North Americans to assume that cities at the east coast of the United states are also east to cities on the West coast of the South American continent.

Another categorical error can arise when judging the distance between places; distances between places situated in different regions tend to be overestimated compared to similar distances between places within the same region (Allen & Kirasic, 1985; Burris & Branscombe, 2005; Carbon & Leder, 2005; Friedman & Montello, 2006; Newcombe & Liben, 1982). Further, the results of both Burris & Branscombe (2005) and Friedman & Montello (2006) suggest, that the perceived distance differences stem from an overestimation of interregional distances, instead of an underestimation of intraregional ones, however the opposite has been found to create the distance mismatch in Carbon & Leder (2005). The distance can also be reflected in route choices, for instance taxi drivers reported factoring in regions in their route decisions (Spiers & Maguire, 2008). In the virtual experiment of Wiener & Mallot (2003), subjects were asked to find the shortest route between a starting point and several goal points. When faced with otherwise equidistant route choices, subjects preferred routes crossing fewer regional borders. Moreover, the effect was not shown by a small number of subjects, who had previously reported that they had not noticed regions, which aligns with the route choice being influenced by the regions.

#### Language can influence spatial reasoning

Information derived from spatial language has been hypothesised to be amodal, given that it may be not be perceptual and given that it may be independent of the input modality. On the contrary, spatial representations are generally seen as modal, at least on a low level of regionalisation. Nevertheless, it is possible to gain functioning spatial information from verbal descriptions alone and studies comparing learning, performance, or errors, point to at least some degree of functional equivalence between information provided by spatial language and spatial representations (Avraamides et al., 2004; Jones et al., 1995; Klatzky et al., 2003; Loomis et al., 2002; Smyth & Scholey, 1996).

However, language does not intrinsically carry spatial information compared to modalities typically employed in orienting, such as vision, hearing or proprioception. And while there is abstraction in spatial representations, leading to more amodal representations, there may be a cost in the conversion of language information to spatial infomation. Fittingly many studies, even when some functional equivalence was found, also found a lower performance when spatial information was given through language compared to more modal information. (Avraamides, 2003; Klatzky et al., 2002, 2003, 2006). Specifically, there was also a lower performance with simple language information compared to virtual sound information, pointing toward a cost specific to language. This is corroborated by the activation of different brain areas for language and visual input of spatial information (Mellet et al., 2002). Interestingly, the studies of Avraamides et al. (2004) and Avraamides & Kelly (2008) found high functional equivalence of spatial language and visual information, either in performance or in subjects showing orientation biases usually connected to physical movement only. However, in both cases subjects showed functional equivalence only after they were induced to form a spatial image out of the linguistic information before the actual testing.

In some cities one may find thematically named adjacent streets, such as a street cluster in New York (Yonkers, NY 10701, United States) where one can get from Maple to Chestnut Street by going through, Oak, Ash, Elm and Linden Street. However, would these streets also form regions within people’s minds? In many instances the streets do not share any stronger functional purpose, oftentimes they might not even share a visual semblance. Often, the street name is not visibly connected to its street at all. Thus, their common denominator would be mostly in the street names alone, something which theoretically can be altered without causing a significant change. Studies have found that visual landmarks are generally preferred over street names; be it in learning of route instructions, following them or giving route instructions to others (A. Tom & Denis, 2003, 2004). Schick et al. (2019) tested whether semantically themed place names could evoke regionalisation as had been found for functionally grouped place names in their experiments. The functional grouping was based on the subunits all being objects typically found in a certain environment (e.g. Campus, canteen, seminar room, lecture hall in a university setting). Semantic grouping on the other hand does not have to imply spatial relatedness, if a semantic group is defined as all items having a commonality on a higher cognitive level.

#### Purpose of the study

Our main goal was to test whether semantically related place names can be used to evoke regionalisation. Thus we wanted to see, whether subjects would prefer routes crossing fewer borders according to the used regionalisation scheme. We used place names standing in an ambiguous relation to each other. To evoke different regionalisations we devised a classification task which would present the place names in slightly different contexts according to the desired classification scheme. If subjects’ route preference reflects the categories they previously formed, it would reinforce the assumption, that the route preference was indeed caused by the semantic groups instead of an inherent other trait in the environment.

## 2 Material and Methods

### 2.1 Subjects

We tested 26 subjects (10 men, 16 women, mean age= 21.27, SD= 4.05). All subjects were able to speak German on the level of a native speaker. Subjects were naïve regarding the purpose of the experiment and testing environment. Subjects gave informed and written consent regarding the experiment and their data and were compensated either with course credit or a payment of 8€ per hour. Two subjects stopped the experiment prematurely due to simulator sickness. 18 out of the total of 26 subjects were tested during the Covid-19 pandemic and thus wore face masks during the entire experiment.

### 2.2 Environment

The environment was created using Unity (Versions 2018.3.0f2 and 20.1). Subjects interacted with the environment using either Oculus Rift goggles (Menlo Park, California 94025, USA) or HTC Vive goggles (HTC Corporation, Taoyuan City 330, Taiwan, (R.O.C)). Both goggles could display 1080 × 1200 pixels per eye at a maximum of 90 Hz, the field of view was 110°. The environment consisted of twelve places with six junctions forming a point-symmetrical hexagon with dead-ends attached to each corner (s. Table 1 and Figure 1 B). The places all looked the same, except that each place had three banners with the place name written on them, which provided the only landmarks (Figure 1 A). To ensure that all places had the same structure, the dead-ends had two blocked roads attached (Figure 1 C), with the blockade being visible from the respective dead-end. It was not possible to look into other places as the junctions were separated by hills which obstructed the view. To move between junctions, subjects had to look in the desired direction and then initiate movement with a button click. The beginning of each path was labelled with the place name of the junction it leads to (Figure 1 B), to facilitate subjects’ orientation.

**Figure 1:**
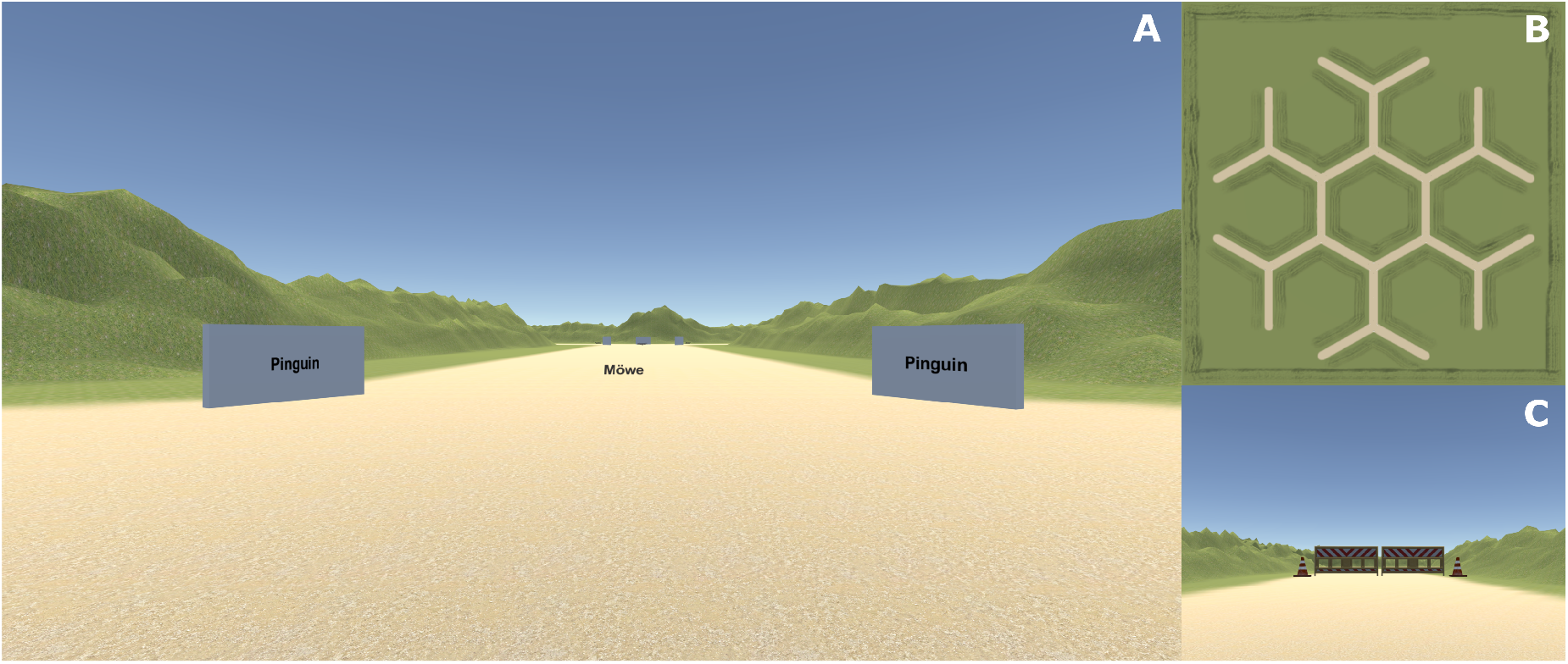
**A**. Each location had three signs displaying the place name only (In the above example “Pinguin”). Each arm was labelled to show the place name of the adjacent location (In the above example “Möwe”). **B**. The experimental environment, all arms are equidistant. Subjects never saw the environment from a birds eye view.**C**. Street block marking at a dead-ends.

### 2.3 Place Names

We used twelve place names, each place name being unique to its respective junction. The place names were chosen so that they stood in an ambiguous semantic relation, meaning that each place name could be grouped into either of two valid categories. Overall two classification schemes were possible each consisting of three categories containing four words each. For example, in Classification Scheme 1, the words ‘Chicken’ and ‘Quail’ would be placed into the ‘Bird’ category alongside ‘Penguin’ and ‘Seagull’, while consistent with Classification Scheme 2, both words could be placed into the ‘Food’ category alongside with ‘Hotdog’ and ‘Hamburger’. Each category consisted of four words (s. Table 1) and subjects could discover one of the two possible overall classifications schemes. Place names that could be grouped into a cohesive group were placed at neighbouring junctions in the testing environment. Thus the environment could be structured into regions in two ways according to the classification scheme used by the subject.

**Table 1:** Place name positions within the environment and the classification schemes they adhere to. Due to the place names ambivalent relations, two equally valid classifications were possible. There were no alternate logical classification schemes found, neither by us nor by the subjects.

| Number | Place Name | Classification Scheme 1 | Classification Scheme 2 |
| --- | --- | --- | --- |
| 1 | Seagull(Möwe) | Birds | Ocean |
| 2 | Penguin (Pinguin) | Birds | Ocean |
| 3 | Quail (Wachtel) | Birds | Food |
| 4 | Chicken (Huhn) | Birds | Food |
| 5 | Hamburger (Hamburger) | USA | Food |
| 6 | Hotdog (Hotdog) | USA | Food |
| 7 | Football(Football) | USA | Sports |
| 8 | Baseball (Baseball) | USA | Sports |
| 9 | Riding (Reiten) | Transportation | Sports |
| 10 | Bicycle (Fahrrad) | Transportation | Sports |
| 11 | Steamboat (Dampfer) | Transportation | Ocean |
| 12 | Ferry (Fähre) | Transportation | Ocean |

### 2.4 Experimental Procedure

The experiment comprised four phases and a post experimental questionnaire. Phases 3 and 4 (Training and Test phase) were conducted in immersive VR.

#### Priming the classification scheme (Classification Phase)

At the beginning of each experiment subjects were asked to do a categorisation task. The twelve place names were presented on a computer screen (30 inch, Acer) and the task was to group the twelve place names into three categories using drag-and-drop (Figure 3). Each category had to consist of four words and had to be named by the subject. Subjects were randomly assigned to one of two groups, which were primed to classification schemes 1 or 2 respectively. Priming was used to increase the likelihood that subjects would form the classification scheme we intended them to use. Two of the category fields were prefilled with one word each, for classification scheme 1, these were ‘Seagull’ and ‘Hamburger’ and for classification scheme 2 ‘Steamboat’ and ‘Hamburger’. These two words therefore had to go into different categories which biased the selection of the overall classification scheme. Other than category size and the prefilled words, there were no further restrictions regarding the categorisation.

**Figure 2:**
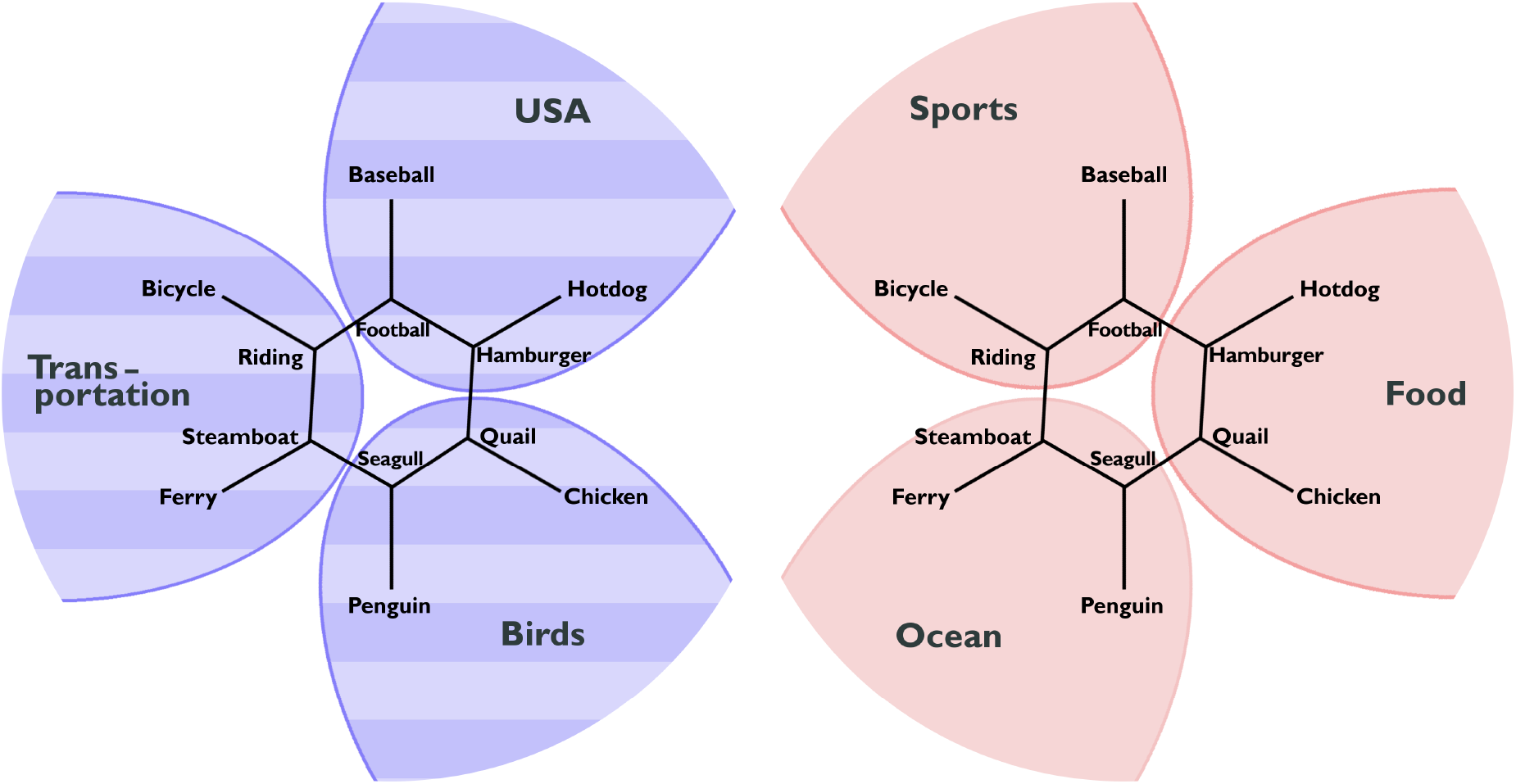
The place names within the environment could be grouped according to either of two equally valid classification schemes

**Figure 3:**
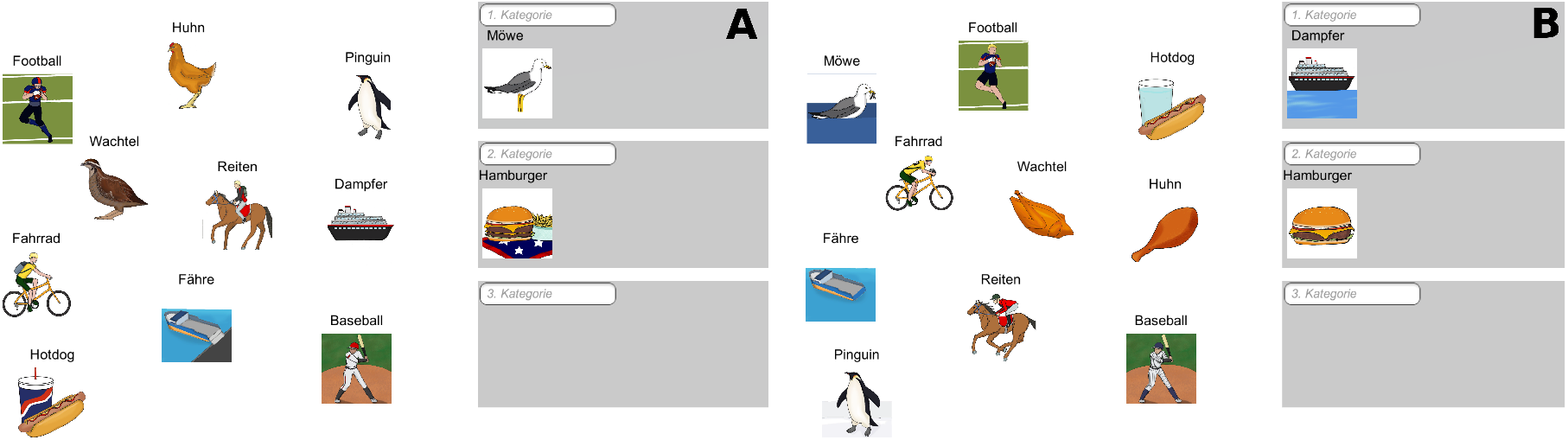
Subjects were to form three categories consisting of 4 words each and to then name the categories. The images accompanying the words and the pre-filled words were changed according to the classification we wanted the subjects to discover. **A**. Shown to induce Classification 1 (Bird, USA, Transportation). **B**. Shown to induce Classification 2 (Ocean, Food, Sports).

Each place name was accompanied by an image showing an example of the denoted object; these images were used in the classification part only, never in the main environment. The images were used to further bias the subjects’ choice of classification scheme, by depicting different, but valid representations of the words depending on the wanted classification scheme. If for example, subjects were primed to classification scheme 1, the words ‘Chicken’ and ‘Quail’ would be accompanied by the images of the respective live animals supporting a “Bird” category. To bias for classification scheme 2, however, the same words appeared alongside images of the processed and cooked meat, supporting classification as ‘Food’.

#### Learning of the environment (Exploration Phase)

In the subsequent exploration phase subjects could explore the main environment in VR which was the same also for the subsequent training and testing phases. There were no requirements for the subject to follow a certain route. The exploration phase ended after the subjects had visited each junction at least once and explored the environment for at least eight minutes. After the exploration phase subjects were asked to sketch the environment on a sheet of paper. The drawings were used to ascertain whether the subjects’ mental image of the environment was accurate.

#### Familiarize subjects with the navigational task (Training phase)

Subjects were transported to a specific junction and given a place name with the task being to the shortest route possible between the starting and goal point (Figure 4). The place names were displayed in the VR goggles as floating words moving with the subjects’ heading and thus always occupied the same position in the field of view. The routes only had one correct solution and spanned three to four segments. Subjects were given feedback on whether they found the shortest route. If the subjects failed, the task would be re-queued, if subjects failed the task four times the route would not get used again. Criterion: How many routes had to be completed?

**Figure 4:**
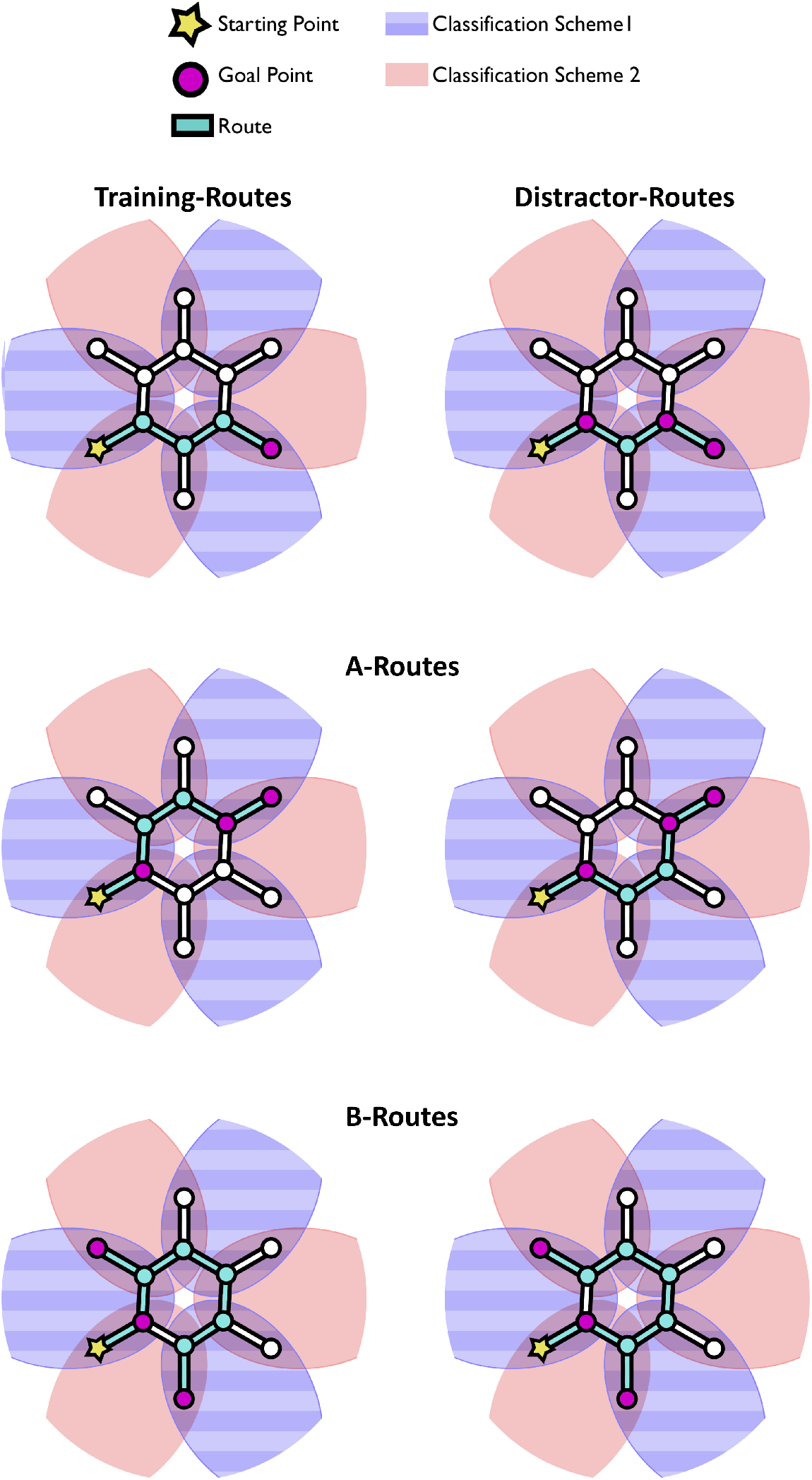
The subject was teleported to the respective starting point at the beginning of each trial. The remaining destinations of each trial were shown to the subject in a steadily varying order. Only the A and B Route types had two possible equidistant route options. One of the route choices in A and B Route types was consistent with the previous classification, while the other was not. As backtracking automatically lead to failure, the first directional choice determines whether the route taken was consistent with the previously formed classification.

#### Testing subjects for their orientation behaviour (Test phase)

The test phase had the same structure as the training phase, the only difference being the route-types given. Unlike in the training phase in which each task had only one shortest solution the test phase now contained tasks with two equidistant routes being being equally short. Further subjects had to find the shortest route between one starting point and three goal points instead of having only one goal. Like in the training phase, subjects had the goal place names displayed before them as a work list To avoid biasing by the order of presentation, the positions of the place names in the work list were randomized every two seconds. To prevent the subject from noticing the existence of equidistant routes, half of the tasks were distractor routes that had only one shortest path. All routes started from a dead end. We used six A-routes, six B-routes and twelve distractor routes (Figure 3). In A-routes, the end goal was the opposite dead-end with intermediary destinations being the corners attached to the respective junctions. There were two possible shortest route options for the A-routes, both spanning five segments, one crossing two regional borders, the other crossing only one. The destinations of the B-routes were the two nearest dead ends to the starting point and the corner opposite to the starting point. Like A-routes B-routes could be solved by either of two shortest routes; B routes spanned nine segments. The destination of distractor routes comprised of the next to nearest dead end either to the left or right of the starting point and the respective corners next to the destination and starting dead ends. Distractor routes only had one solution and spanned four segments.

#### Asking about personal information and feedback (Post Experiment Questionnaire)

We used the post experiment questionnaire to access the subjects’ individual sense of difficulty and content toward the classification part. Further, subjects were questioned on whether they thought that the classification part may have influenced them during orientation. We also recorded information relating the subjects as previous studies regarding virtual reality orientation have found that women and non-video gamers are more susceptible to simulator sickness, which in turn can affect performance (Weech et al.,2020; Grassini et al., 2021).

#### Data Analysis

The effect of classification consistent route choices (CCRC) consists in a preference for routes crossing a lower number of region boundaries over equidistant routes which cross a larger number of region boundaries. In our experiment, we test for routes avoiding the previous classification scheme’s borders, as any route choice will be consistent with one categorization scheme or the other, as regionalisation was ambiguous. Indeed, we can use the subject’s route choices to infer the underlying regionalisation used in each trial. For this, we consider the region, from either of the two categorization schemes, which contains the first three places of a given route, i.e. the starting point in a dead end and the two next places on the ring. It is determined after the first directional decision (left or right) made by the subject. We then call the route chosen in the respective trial to be consistent with the regionalisation containing this region. E.g. a route starting with the places penguin-seagull-quail would be called consistent with categorization scheme 1, while a route starting with the places penguin-seagull-steamboat would be called consistent with categorization scheme 2. This analysis can be applied to both route types, A and B.

## 3 Results

### 3.1 Classification Tasks

In the following we will look at route choices separately for trials in which the subject was able to find the shortest route and trials where the subject was not able to do so. This was done as it is questionable whether the route choice a subject made can reflect the tendency for CCRCs truthfully if the person was possibly disoriented. All routes in all conditions were solved in less than four tries, except for one B-route type for one subject which was excluded from further analysis. The classes chosen in the classification phase were checked for every subject. The data of four subjects who formed classes with unexpected item combinations were excluded, as the resulting regions would have been disconnected. Further, at least one of the categories in these cases was either illogical or a compound class, which may have weakened the category and the resulting regionalisation (e.g. a ‘Flying’ category which consisted of ‘Penguin’, ‘Football’, ‘Baseball’ and ‘Seagull’ or a ‘Food and Means of Movement’ category). There was no case of a subject forming categories with unexpected item combinations where all three classes were also logical. One data set was excluded even though the subject formed categories with the expected content, as the class a name suggested that the subject formed the class based on illogical criteria (the subject formed an ‘Object’ category out of ‘Penguin’, ‘Ferry’, ‘Steamboat’, ‘Seagull’). Further two subjects dropped out, which resulted in 19 included subjects.

Out of the usable datasets, ten subjects chose Classification 1 and nine subjects chose Classification 2. All of those subjects chose the classification type in accordance to our priming in the classification phase. All subjects gave either the highest or second highest rating regarding their satisfaction toward their formed categories in the classification phase (1(very content) – 5(discontent), mean rating for all subjects = 1.583, mean rating of included subjects = 1.474). Further, all subjects rated the classification task’s difficulty as low to medium (1(easy) – 3(hard), mean rating of all subject= 1.375, mean rating of included subjects = 1.210). Out of the 19 subjects 12 stated that they felt their classification had influenced their route decision. However, there was no significant difference in making CCRCs between subjects who felt influenced (mean percentage of CCRCs= 65.28%, SD =18.33) and subjects who did not (mean percentage of CCRCs= 56.9%, SD =20.1, t(36)=1.312, p= 0.629, two sample t-test).

#### Classification consistent route choices (CCRC)

There were no differences in the percentage of CCRCs between A- and B routes (mean CCRC percentage for A-routes= 62.3%, SD= 17.43; mean B-routes= 62.1%, SD= 21.23, t(36)=0.056, p=0.978, two sample t-test). We therefore merged the datasets for both routes for further analysis. We tested for correct trials whether the subjects’ chosen classification influenced their route choices, thus whether they showed the CCRC effect with the regionalisation previously discovered in the catergorization phase of the experiment; the data appears in Figure 5. A *χ*^2^-test for categorical data revealed a significant CCRC effect (Figure 5, *χ*^2^(1, N= 227)= 13.86, p*<*0.001). The CCRC preference was still visible when the choices of failed trials were added, thus when the effect was looked at for all trials (*χ*^2^-test for categorical data, *χ*^2^(1, N= 260) = 13.34, p*<*0.001). It should be noted though, that the number of failed trials was much smaller than the number of successful ones. Subjects who performed poorer in the test phase in finding the shortest route did not show a lower preference for making CCRCs (r=-0.230, p= 0.345, Pearson’s correlation coefficient).

**Figure 5:**
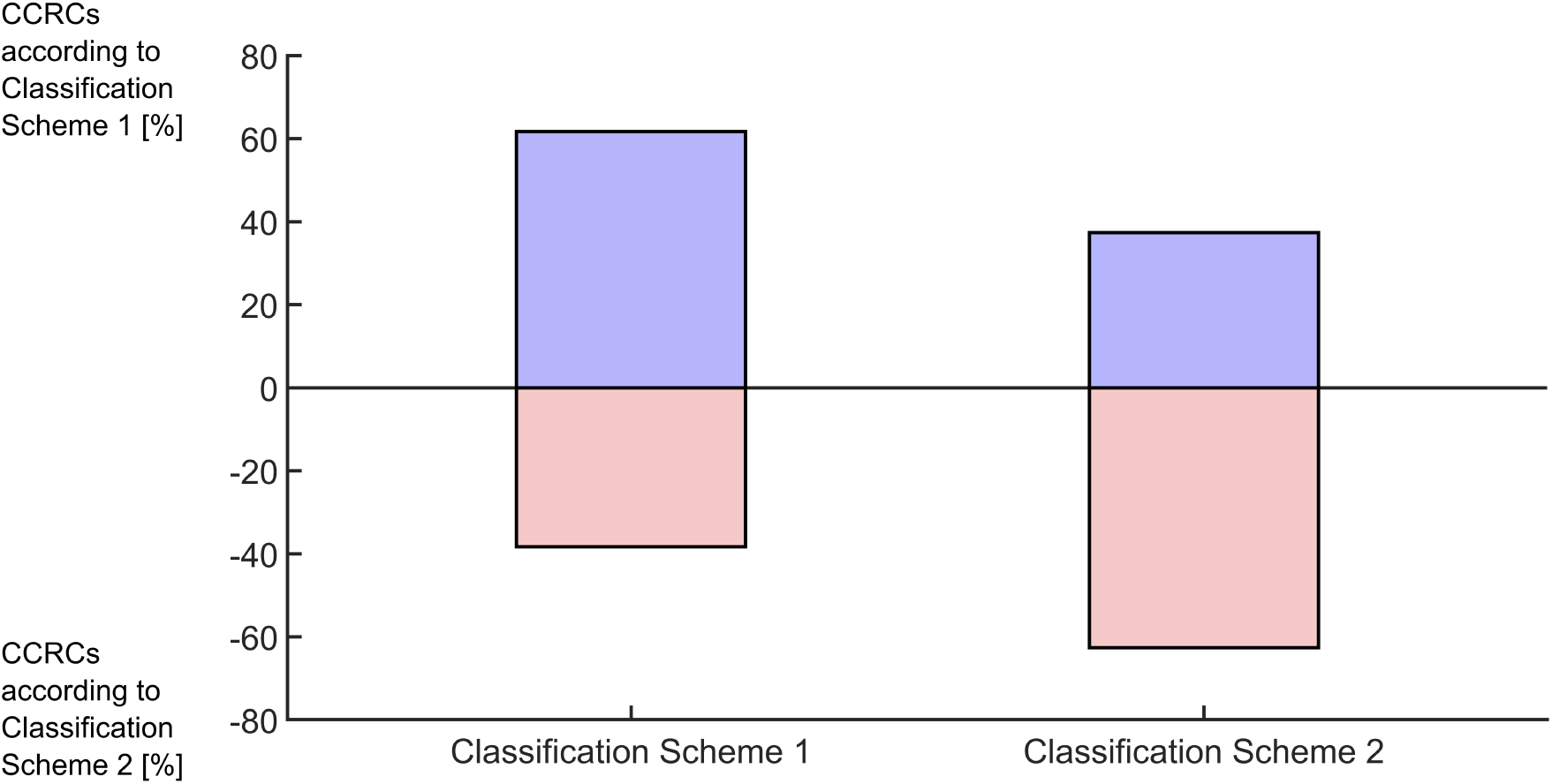
Subjects preferred routes with fewer border crossings according to the discovered classification scheme. For trials in which the subject previously discovered the classification scheme 1 the absolute number of CCRCs was 74 compared to 46 non CCRCs. For trials in which the subject previously discovered the classification scheme 2 the absolute number of CCRCs was 67 compared to 40 non CCRCs.

#### Directional Biases

We tested the direction of all first choices the subjects made in either A or B routes for each trial, regardless whether they found the shortest route or not. We found that subjects displayed no general bias either to the left or right for either route type (A-routes p(115)= 0.307, B-routes p(143)= 0.080, Sign test). We also found no directional bias when looking at all choices for all route types (p(499)= 0.823, Sign test).

#### Route precision

Subject made more errors in B-routes than in A- or distractor routes, and thus subsequently needed more trials to complete all routes in the respective condition ((F(2,36)=13.85, p*<*0.001). To complete all six A-routes subjects needed 6.105 ± 0.315 tries, to complete all six B-routes subjects needed 7.579 ± 1.953 tries, to complete all twelve distractor-routes subjects needed = 12.737 ± 1.447 tries). There was no difference in errors between the A-routes and distractor-routes, however there was a difference between B-routes and both A-routes and distractor-routes (p(AB)*<*0.001, p(AD)=0.522, p(BD)*<*0.001).

#### Navigation Duration

We calculated the time per segment, i.e. the average speed, considering only successful routes. The time per segment the subjects took was significantly different for all route types F(2, N=113)= 27.97,(p*<*0.001). Subjects took longest for the B-routes (M = 6.78 seconds, SD=2.63) followed by A-routes (M = 6.09 seconds, SD= 2.24) and shortest for the distractor routes (M = 5.41 seconds, SD= 1.63). There was a significant interaction of subjects and route type (F(36, N=19)= 3.24, p*<*0.001).

#### Individual Analysis

The number of subjects who made more CCRCs than non-CCRCs was significantly higher than the number of subjects who did not (see Figure 6, p(18)= 0.013, Sign Test). Out of 19 subjects 14 made CCRCs in more than 50% of all choices, with the meridian at 66.7% (SD= 13.96) There were no sex differences regarding the number of trials needed for either route type (mean number of trials men needed for all A-routes= 6.00, SD=0; trials women needed for all A-routes= 6.13, SD=0.35, t(17)=0.742, p= 0.468; mean number of trials men needed for all B-routes= 7.53, SD=2.22; trials women needed for all B-routes= 7.53, SD=1.96, t(17)=-0.192, p= 0.850). There further was no difference in the tendency to make CCRCs for men (mean percentage of CCRCs made= 59.25%, SD= 15.99) and women (mean percentage of CCRCs made= 62.42%, SD= 16.77, t(36)= 0.479, p= 0.635). Likewise, video gaming experience was neither correlated to route precision in A-routes (r(17)= 0.295, p= 0.221) nor in B-routes (r(17)= 0.303, p= 0.208, Pearson’s correlation coefficient). Further there was no correlation of gaming experience and inclination for CCRCs (r(36)= 0.141, p= 0.399, Pearson’s correlation coefficient).

**Figure 6:**
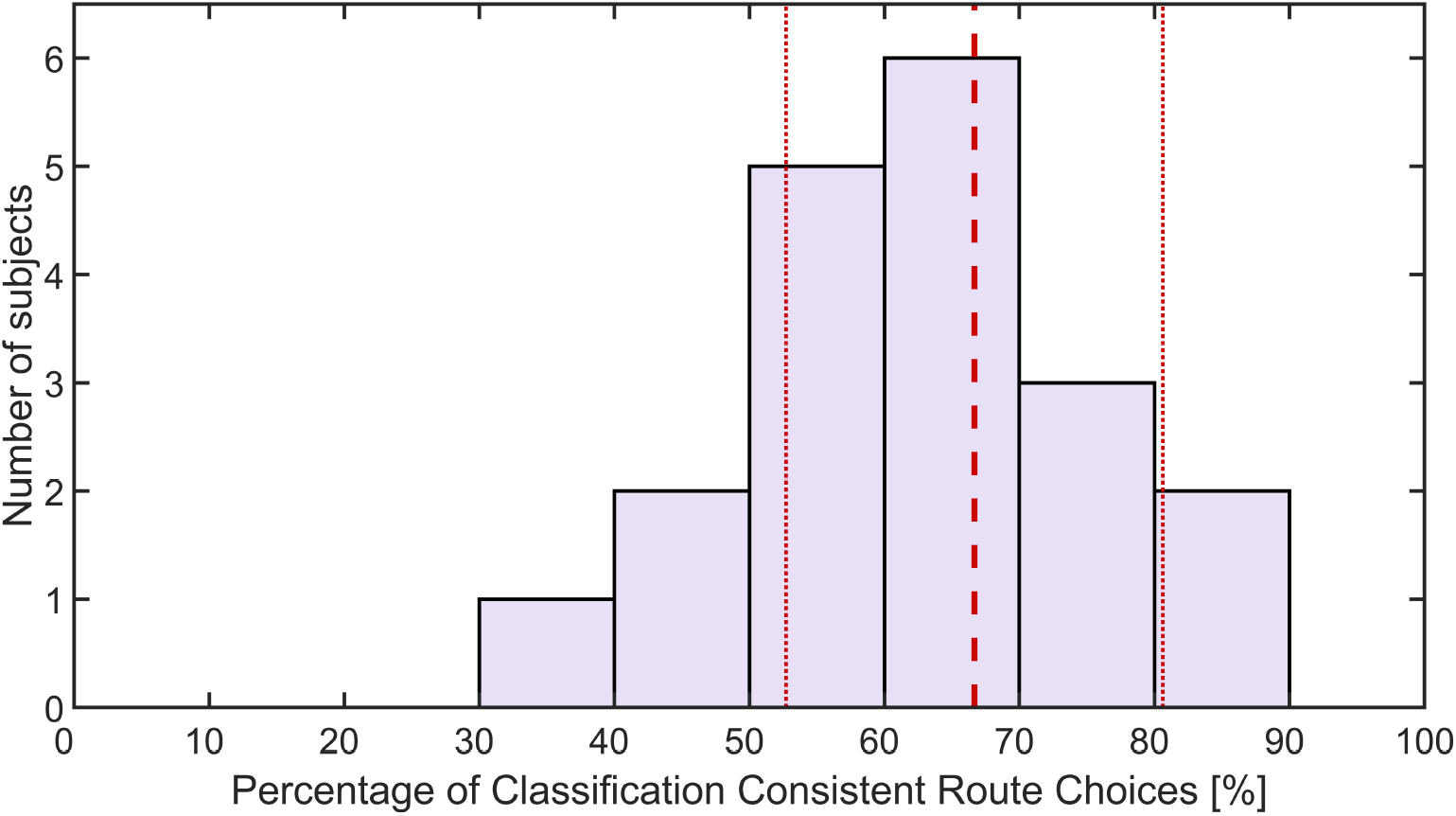
The number of subjects according to the subjects’ adherence to making CCRCs. The thicker dashed line represents the median while the dotted lines signify the standard deviation. While subjects differed in their orientation behaviour, the majority of subjects did prefer routes with less border crossings.

## 4 Discussion

The study examined whether the perceived semantic relation of place names can induce a regionalisation effect when semantic relations are ambiguous. Indeed, subjects avoided crossing the semantic borders of their previously discovered place name groups, indicating that regionalised route planning depends on perceived semantic grouping. The simple hexagonal structure of the maze rules out the definition of regions by the density of connections and puts the focus on the role of similarities between places and landmarks. Wiener & Mallot (2003) show that semantic relations qualify as similarities in this sense, i.e that they do induce regionalisations. However, the results by Schick et al. (2019) indicate, that semantic relations do not always lead to regionalisation and that functional relations may be more important than similarities of kind. The present study provides clear evidence that region formation is not just determined by a fixed membership of place names to semantic classes. Rather, the co-activation of objects used as place descriptors in a preceding task influences regionalisation. This ties in with prior findings, that spatial perception can be influenced by previously established connotations that may be different for different observers (Burris & Branscombe, 2005; Carbon & Leder, 2005; Friedman & Montello, 2006). The CCRC effect is slightly less than the region dependent route choice effect in Wiener & Mallot (2003). Given that the place names were chosen to equally fit into two categories, they might be less prototypical for either category compared to place names without such a constraint (Rosch & Mervis, 1975). As reduced prototypicality may have reduced the category and regionalisation strength, a reduced effect was therefore to be expected.

The role of semantically related place names was also studied by Schick et al. (2019) who found that using textual names instead of visual objects can weaken the induction of regions by semantic relations to the point of insignificance. We suspect two main reasons why the effect of semantically induced regions was strong enough to induce a significant orientation bias in our experiment: decreased orientation difficulty and the visually supported classification task preceding the experiment.

Firstly, our experiment was more easily navigable due to the paths being labelled with the name of the destination the path lead to. In addition to easier navigation, the path labelling may have also facilitated learning the environment. Also, problems of teleportation. i.e. the disorientation caused by sudden scene change (Bolte et al., 2011; Bowman et al., 1997; Cherep et al., 2020), would be reduced. Having more information regarding their position, after having been teleported at the beginning of each trial, subjects may have had less difficulty reorienting themselves. This is corroborated by our experiment having a lower route failure rate and only one performance dependent route omission.

Secondly, the classification phase of our experiment may have put the initially unrelated place names into a functional context. Indeed, such functional contexts did support regionalisation in the Schick et al. (2019) study. For example, functional relations between place names such as “library” and “lecture hall” as opposed to “barn” and “stable” did induce region dependent route choice and even led to the report of little narratives about visiting a university campus or respectively a farm by some subjects. The classification task also provided visual images of the place names which may have supported regionalisation. Pantelides et al. (2016) found that people can integrate two spatial frameworks, a verbally learned and a visually learned, into one. Yet if the integration is difficult, e.g. because of different learning perspectives, the integration does not happen automatically, but may still occur later on during retrieval. It has been found in A. Tom & Denis (2003, 2004) that landmarks are generally more important and readily used in spatial learning and wayfinding than street names or descriptions. However, A. C. Tom & Tversky (2012) also found that vividly described streets can be remembered even better than landmarks and that recall was better in people with stronger mental imagery abilities. Moreover, it has been found that vividness, a stronger mental image, can enhance recall in general (Collins et al., 1988). Therefore, the images in the classification phase may have eased forming a mental image which may in turn have enhanced recall and thus furthered regionalisation.

Finally, drawing the environment may have strengthened the regionalisation the subjects had or even created it. Subjects may have used their own drawings to form a mental map. It has been found that access to prior information, especially in the form of a map can facilitate adopting a different view than the immediate egocentric one (Meneghetti & Pazzaglia, 2021), which would presumably strengthen the formation of regions as well.

We had four subjects who were excluded from analysis as they had formed at least one illogical or inconsistent category. It is not clear how they formed the illogical categories; one possibility would be that the subjects settled on inconsistent categories because they did not find a way to form sensible categories out the place names. However, no subject rated the classification task as hard and all subjects gave either the highest or second highest satisfaction rating regarding their formed categories. This would suggest that the subjects considered their categories logical.

We tested a possible correlation between contentment with the formed categories and likelihood for CCRCs as higher contentment would presumably be tied to a stronger regionalisation. There was no significant correlation between contentment in the formed categories and likelihood for CCRCs. However, given the overall high satisfaction ratings, we suspect a ceiling effect. Thus, based on the data alone the study can neither support nor deny a positive correlation between the contentment and strength regarding the classes.

The place names used in this experiment have been carefully selected to allow the ambiguous categorization required for the design. The high satisfaction of the subjects with their classification seems to support this selection. Moreover, with ten subjects choosing Classification 1 and nine choosing Classification 2, both classifications seem to be of similar strength. The balance in both classifications is further supported by the fact that all subjects who formed logical and consistent categories also made the classification suggested by the classification task.

It is not clear whether subjects’ first directional decision was made on a local or on a global scale, i.e. whether subjects chose the direction that allowed them to stay in the current semantic category, or if subjects chose the route with the least number of border crossings. While our question whether subjects develop and act upon regions is answered by either case, it may be of interest to discern between a global and local effect. However, this would need a more complex and possibly asymmetrical testing environment.

In our experiment the control hierarchy that determines the preferred routes was influenced by the representational hierarchy, reflected in the grouping of place names. Given that subjects readily accepted both classification schemes, there is the possibility that humans may have multiple parallel representations for a single environment. These parallel hierarchies can activate separately, with each leading to a different spatial representation of the environment. The current context can then cause one hierarchy to be activated over the other. As we exposed each subject to one classification scheme only, our results cannot confirm the theory, however an experiment in which subjects are primed to switch from one regionalisation to another might be of interest in further studies.

In conclusion, semantic relations in place names are indeed sufficient to evoke regionalisations. By imposing a cost on region crossings, the regionalisation can then influence orientation behaviour in turn. By changing the context of the place names, we could influence the semantic relations the subjects would focus on. Thus we could nudge subjects into forming one regionalisation scheme over another. Our results thus indicate, that a single environment can support multiple regionalisations, which can lead to differing orientation behaviours.

## 5 Compliance with Ethical Standarts

### Disclosure if potential conflicts of interests

The authors declare no conflict of interest.

### Informed consent

Informed written consent was obtained from all individual participants included in the study.

### Research involving human participants and/or animals

The manuscript does not contain data from clinical studies, patients, or non-human animal data.

### Data availability statement

Data is available at request from the authors.

### Author contribution

All authors contributed to the study conception and design. L.L. performed the experiments and analysed the data. All authors jointly prepared the manuscript. All authors read and approved the final manuscript.

### Funding

The research reported in this paper was carried out at the Department of Biology of the University of Tübingen. Additional support was provided by the Deutsche Forschungsgemeinschaft (DFG - German Research Foundation), grant no. 381713393, within the Research Unit FOR2718: Modal and Amodal Cognition.

## Acknowledgement

We are grateful to Svenja Zehender, Leonie Mödl and Jule Wildt for their contributions to the development of this experiment

